# Are Solid Particles Ready for Prime-Time Proteomics?

**DOI:** 10.1101/2025.05.22.655475

**Authors:** Eduardo S. Kitano, Yana Demyanenko, Shabaz Mohammed

## Abstract

We evaluate the performance of nonporous C-18 stationary phases in high-speed proteomics workflows. We employed two commercially available sub-2 µm nonporous particle (NPP) materials, ODS-IIIE (1.5 µm) and SOLAD (1.0 µm), to fabricate analytical columns using 150 µm internal diameter (i.d.) fused silica capillaries to ensure compatibility with nano-UHPLC and nano-HPLC pressure regimes. Using ‘long’ NPP columns (15 – 25 cm) connected to a conventional nano-UHPLC, we found that both materials supported efficient peptide separations within the flow rate and pressure ranges typical of nano-UHPLC systems. Shorter columns, used with the 10- and 16-minute Whisper Zoom 120 and Zoom 80 methods on the Evosep One HPLC, demonstrated competitive separation performance compared to columns packed with fully porous (FPP) and superficially porous particles (SPP), achieving full-width at half maximum (FWHM) values below 2 seconds (Zoom 120) and 3 seconds (Zoom 80). DIA LC-MS/MS analysis of the same digest demonstrated that NPP columns provided comparable or superior performance relative to FPP and SPP materials. These findings establish NPP-based columns as a viable and competitive alternative to FPP and SPP materials, particularly suited for high-throughput and high-sensitivity proteomics applications.

## Introduction

Recent technological advancements in mass spectrometry (MS) have significantly increased sensitivity and acquisition speeds^1, 2^, reshaping the design and optimization of liquid chromatography–mass spectrometry (LC-MS) workflows.^3–6^ As a result, short-gradient LC-MS/MS methods, particularly when integrated with data-independent acquisition (DIA) strategies, are increasingly employed in high-throughput applications such as clinical proteomics^7, 8^, biomarker discovery ^9, 10^ and single-cell proteomics.^11–13^

Underlying much of this progress is the widespread adoption of sub-2 µm fully porous C-18 particles (FPPs) in column design, which have proven crucial in proteomics due to their high surface area, excellent mass transfer properties, and broad compatibility with both nano- and micro-flow systems.^14, 15^ While FPPs remain the dominant stationary phase, alternative technologies like superficially porous particles (SPPs) and nonporous particles (NPPs) have shown considerable potential for LC-MS/MS applications.^16–18^ Both SPP and NPP C-18 phases offer potentially distinct advantages over FPPs, including superior mass transfer kinetics, reduced peak dispersion, and higher resolution at high flow rates – features that are particularly advantageous for high-speed separations.^19–22^ Due to their inherently low surface area, nonporous particles have limited sample-loading capacity restricting their application to low sample amounts^23^ while the higher surface area of the porous shell in SPPs enables loading capacities comparable to those of FPPs.^24^ The enhanced chromatographic performance of these materials results in sharper peaks^23^ and stronger signal intensity, which in data-dependent acquisition (DDA) improves precursor ion selection, and in DIA enhances the separation of co-eluting species, leading to cleaner MS/MS spectra and more confident peptide identification and quantification.^25, 26^ The potential higher-separation power of these materials position both SPP and NPP as attractive tools for the next generation of fast and high-sensitive proteomic workflows.

Although the use of NPP-based columns in proteomics has been limited^25^, the sub-nanogram sample requirements of modern mass spectrometers now position them as a compelling option for high-efficiency separations in low-input proteomic applications. In this study, we investigate the applicability of NPP and SPP columns for high-throughput proteomics workflows. Separation performance was evaluated using the two newly established short-gradient methods on an Evosep One LC system and compared with established C-18 materials employed in peptide separation. Coupled with DIA, we compared the peptide and protein identification efficiency of short NPP and SPP columns to that of FPP columns using a human cell lysate.

## Experimental section

Detailed methods are provided in the Supporting Information.

### Protein Digestion and Sample Preparation

Bovine serum albumin (BSA) and α-casein proteins (Sigma), and Expi293F cell (Gibco) lysate were submitted to in-solution LysC/trypsin digestion as previously described.^27^ The resulting peptide samples were resuspended in either 5% formic acid / 5% DMSO for nanoUHPLC analysis or in 5% formic acid / 0.015% n-Dodecyl-β-D-maltoside (DDM) prior to loading onto Evotip pure (Evosep Biosystems) for subsequent nanoHPLC analysis.

### Column Packing

Nonporous, fully porous, and superficially porous C-18 materials were prepared as slurries at a concentration of 100 mg/mL. Specifically, Reprosil Gold™ (fully porous), ODS-IIIE™ (nonporous), and SOLAD™ (nonporous) were slurried in 100% acetone, while Luna Omega Polar™ (fully porous) and Kinetex® (superficially porous) were slurried in 100% methanol. Fused silica capillaries (Polymicro Technologies) with internal diameters (i.d.) of 75, 100, and 150 µm were pulled using a P-2000 laser puller (Sutter Instrument). These capillaries were then packed with the respective C-18 materials using a PC8500 Pressure Injection Cell (Next Advance), following the protocol described by Kovalchuck^28^, with the production of a self-assembled particles as a frit at the outlet of the columns.^29^ Seventy-five micrometre i.d. capillaries were packed with fully porous Reprosil Gold™ (1.9 µm, Dr. Maisch) and Luna Omega Polar™ (1.6 µm, Phenomenex); a 100 µm i.d. capillary with superficially porous Kinetex® (1.7 µm, Phenomenex); and 150 µm i.d. capillaries with nonporous ODS-IIIE™ (1.5 µm, Eprogen/Promigen Life Sciences) and SOLAD™ (1.0 µm, Glantreo). Following packing, the column beds were consolidated by applying a constant flow of 70% acetonitrile at 800 bar for 2 hours. Columns were then cut to the desired length, and a sol-gel frit was created at the outlet according to the method described by Maiolica.^30^ Figure 1 schematically illustrates the different C-18 materials and the columns fabricated in this study.

**Figure 1.**
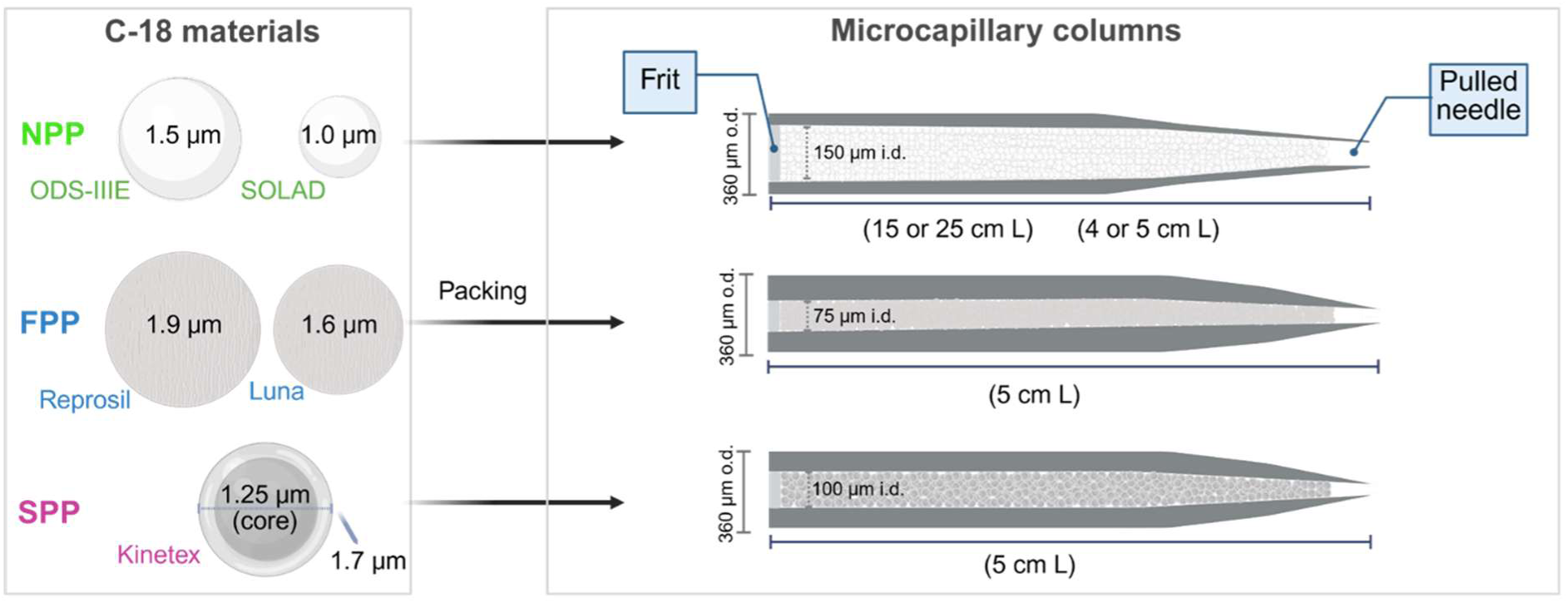
Schematic representation of the C-18 stationary phases and column configurations employed in this study. The left panel presents the morphological characteristics of nonporous (NPP), fully porous (FPP), and superficially porous (SPP) C-18 particles, with particle sizes represented proportionally. The right panel summarizes the key specifications of the columns packed with each respective particle type.

### Nano-Liquid Chromatography and Mass Spectrometry

To optimize flow rates and column temperatures for the NPP C-18 columns, 10 fmol BSA/casein digest was separated using an Ultimate 3000 RSLCnano system (Thermo Fisher Scientific) separated on a 25-cm ODS-IIIE and a 15-cm SOLAD analytical columns. The chromatographic separations were performed across a temperature range of 30 to 70 °C (at a constant flow rate of 200 nL/min), and at flow rates ranging from 50 to 500 nL/min at 40 °C (ODS-IIIE) and 50 °C (SOLAD), utilizing a 15-minute linear gradient of 7-55% solvent B (5% DMSO, 0.1% formic acid in acetonitrile). The BSA/casein digest was also subjected to a 15-minute separation using shorter ODS-IIIE (5 cm) and SOLAD (4 cm) columns at 200 nL/min at 20 °C. To assess and compare the separation performance of NPP columns with that of FPP and SPP columns, a 10 fmol BSA/casein digest was analysed using the predefined Whisper Zoom 120 (10-minute gradient) and Zoom 80 (16-minute gradient) methods on an Evosep One HPLC system (Evosep Biosystems). Increasing amounts of Expi 293F digest (0.25 to 50 ng) separated by the Whisper Zoom 120 and Zoom 80 methods was used for the assessment of peptide loading capacity of 5-cm ODS-IIIE and 4-cm SOLAD columns. Data-Dependent Acquisition (DDA) mass spectrometry analyses were performed on a Q Exactive mass spectrometer (Thermo Fisher Scientific).

Twenty nanograms of Expi 293F digest was separated on NPP, FPP, and SPP C-18 columns using the Whisper Zoom 120 and Zoom 80 methods of the Evosep One coupled to an Orbitrap Exploris 480 mass spectrometer (Thermo Fisher Scientific) operating in Data-Independent Acquisition (DIA) mode.

### Raw Data Analysis

Full-width at half maximum (FWHM) and peptide intensities were manually calculated/obtained from the extracted ion chromatograms (XICs) of selected peptides from BSA/casein or Expi 293F digests using Freestyle™ software (Thermo Fisher Scientific). Data were further processed using GraphpadPrism software. DIA LC-MS/MS data were analysed by DIA-NN software^31^ using library-free search. Spectral library was predicted in silico from the Uniprot human proteome database (Proteome ID: UP000005640, downloaded in August 2022, 79,759 sequences). Methionine oxidation and cysteine carbamidomethylation were set as variable and fixed modification, respectively. Enzyme specificity was set to trypsin/P, up to one missed cleavage, and minimum peptide length of 7 amino acids was allowed per peptide. Precursor FDR was estimated and filtered to 1%. The LC and MS method details are described in Supporting Information. Raw files and results from this study have been deposited to ProteomeXchange Consortium via the PRIDE^32^ partner repository with the data set identifier PXD064019.

## Results and Discussion

### Evaluation and Optimization of Nonporous C-18 Columns for NanoLC Peptide Separation

MS sequencing speed improvements have created the need to generate high peak capacity short LC gradients with equally short loading and regeneration times. Such criteria require shorter columns than those typically utilised in proteomics, where longer columns and longer gradient are *de rigueur*.^33, 34^ For such experiments 5 cm columns are the common length as found on Evosep one systems. We wanted to explore the role of particle type and its size at such column lengths. The use of smaller particles in chromatographic separations results in higher column efficiency^20^, which can further reduce column length to minimize dead volume and potentially improve peak capacity. However, smaller particles also lead to the increase in backpressure, which might require the use of ultrahigh-pressure LC (UHPLC)^35^ or shorter than desired columns to manage the increased diffusional resistance. Given the favourable mass transfer kinetics^21^ and high chromatographic efficiency of sub-2 µm nonporous C-18 particles across a broad range of linear velocities^36^, we hypothesize that these materials are promising candidates for the development of rapid and highly sensitive proteomics methods.

In this study, we chose two commercially available nonporous C-18 materials: ODS-IIIE™ and SOLAD™, with a particle diameter of 1.5 µm and 1.0 µm, respectively (Fig. 1). The smaller particle size of SOLAD is expected to enhance efficiency, but also to increase column backpressure. Preliminary experiments indicated that the relationship between optimal linear velocity and flow rate differs between nonporous and porous particles. Typical flow rates of nanoLC systems are in the 100-400 nL/min range. To ensure compatibility with nanoLC while evaluating performance characteristics specific to nonporous particle (NPP) columns, we selected a 150 µm internal diameter (i.d.) capillary as the starting format. Using this configuration, we prepared 15 cm and 25 cm columns packed with SOLAD and ODS-IIIE stationary phases, respectively. A 15-minute gradient was applied using a BSA and casein digest standard to assess optimal flow rate, temperature, and the associated pressure profiles. The separation window and FWHM values of selected peptides were used as a metric for chromatography performance. As expected, system pressure increases linearly with the increase in flow rate at constant temperature (Fig. 2A, Table S1). Even with the reduction in the column length and higher temperature, SOLAD generated higher backpressures than a significantly longer ODS-IIIE column (15 cm versus 25 cm) at flow rates above 150 nL/min. To mitigate the increased pressure associated with higher flow rates, the column temperature was set to 75 °C for flow rates exceeding 350 nL/min (SOLAD) and 400 nL/min (ODS-IIIE). Under these conditions, we observed a compression of the elution window and a loss of resolution, consistent with the reduced residence time of the peptides due to the higher mobile phase volume passing through the column per unit time (Fig. S1 and S2). Good separation was observed between 150 nL/min and 350 nL/min. We chose to focus on 200 nL/min which is the typical flow rate used for nanoLC-MS. Across the tested temperature range, the ODS-IIIE column exhibited lower FWHM values than the SOLAD column (Fig. 2B, Table S1). For the ODS-IIIE column, FWHM remained relatively consistent, ranging from 2.3 to 2.5 seconds, with the lowest value observed at 40 °C. In contrast, the SOLAD column exhibited greater variability in FWHM (3.7 to 4.6 s) across the same temperature range, suggesting a more pronounced temperature effect on peak width compared to the ODS-IIIE column. The SOLAD column’s lowest FWHM was observed at 50 °C (3.7 s). Under optimal conditions, ODS-IIIE and SOLAD showed similar chromatographic profiles, and FWHM of 2.3 s and 3.7 s, respectively (Fig. 2C-D). These findings support the use of a 150 µm i.d. column as a suitable option for implementation in nanoLC systems, offering compatibility with the pressure constraints at typical flow rate regimes.

**Figure 2.**
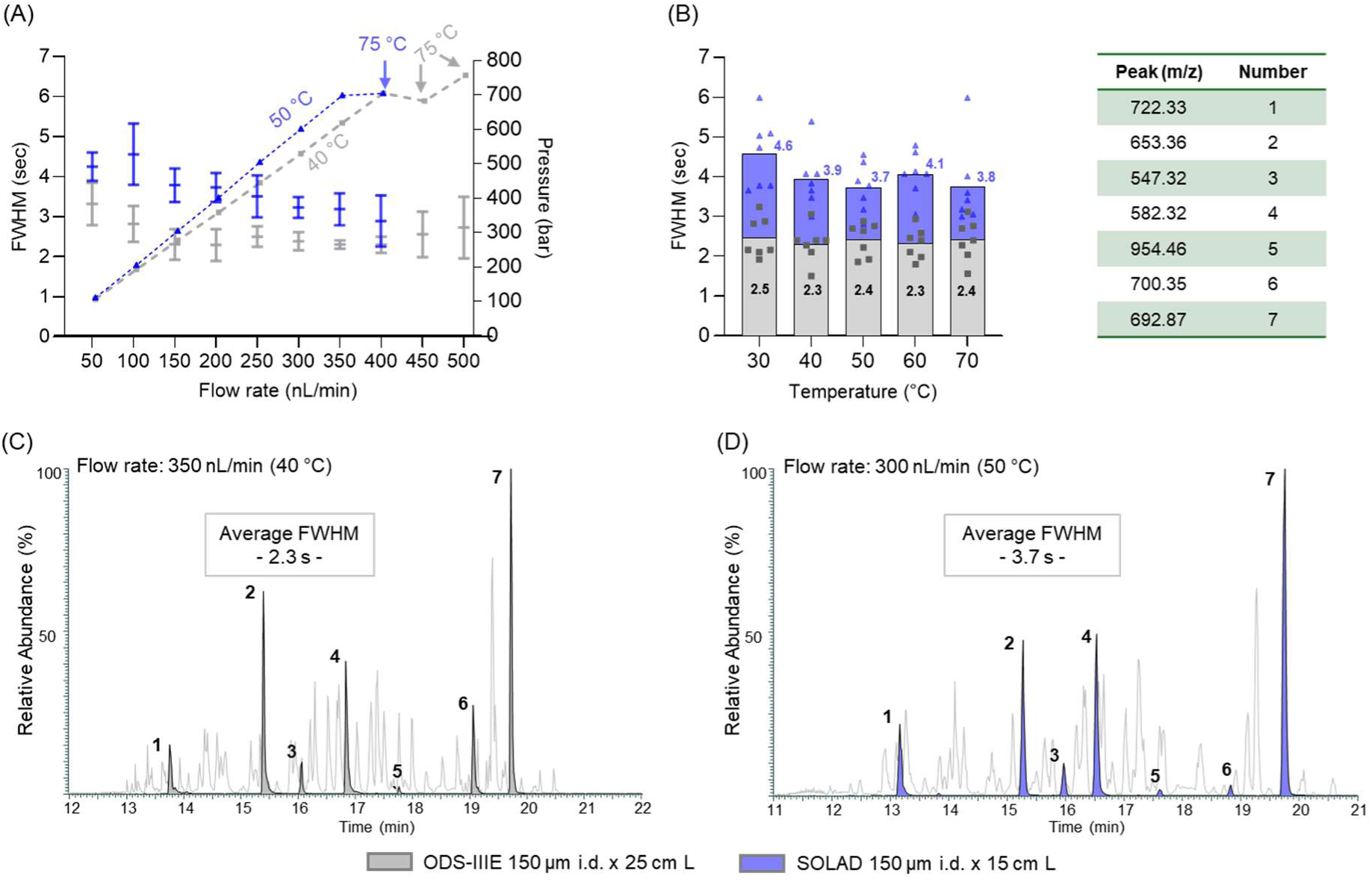
Optimization of non-porous particle columns for peptide separation. Average full-width at half maximum (FWHM) of seven peptides from a 10 fmol bovine serum albumin/α-casein tryptic digest, separated using a 15-minute gradient, was assessed across varying flow rates (A) using 150 µm i.d. × 25 cm ODS-IIIE and 15 cm SOLAD columns, and across varying column temperatures (B). Error bars indicate standard deviation (n=3). Representative base-peak (BPCs) and extracted ion chromatograms (XICs) under optimized conditions are displayed for the ODS-IIIE (C) and SOLAD (D) columns. Peptide peaks are represented by the mass-to-charge ratio (m/z) shown in the table. LC-MS/MS analysis was performed on an Ultimate 3000 RSLCnano system coupled to a Q Exactive mass spectrometer.

### Short Nonporous C-18 for Fast Liquid-Chromatography

We hypothesized the Evosep One with its newly developed Whisper Zoom methods operating at 200 nL/min could be a suitable platform for the evaluation of NPP columns in fast LC-MS/MS. To ensure compatibility with the predefined methods and the pressure limits of the HPLC system, we chose a 5 cm column packed with ODS-IIIE particles, and, given the higher backpressure associated with the smaller SOLAD particles, a shorter 4 cm column was selected. First, using an Ultimate LC, we conducted 15-min LC-MS/MS analyses of a BSA and casein digest at 200 nL/min using both long and short ODS-IIIE and SOLAD columns (Fig. S3). Compared to longer columns, the shorter ODS-IIIE and SOLAD columns exhibited, as expected, significantly broader peaks and increased tailing across the entire elution range. This included early-eluting, known BSA hydrophilic peptides with mass-to-charge ratio (m/z) 488.54, 625.78 and 722.33, which can be used as markers to assess the separation of hydrophilic species. This phenomenon is likely attributable to the mismatch in analyte retentivity between the trap and the analytical columns, which induces band broadening^25, 37^ that is exacerbated by reduced column efficiency, such as that caused by shortening the analytical column. In the Evosep system, elution from the trap (Evotip) is diluted prior to reaching the analytical column, enabling reconcentration^3^ and thereby minimizing the impact of any selectivity mismatch. Next, we assessed the performance of the two short NPP columns on the Evosep system using the Whisper Zoom 120 (10-minute) and Zoom 80 (16-minute) methods, employing the same standard sample (Fig. 3). A significant reduction in peak tailing and broadening was observed for both NPP columns under both methods. This suggests that the selectivity mismatch between the trap and analytical columns was indeed a primary cause of the previously observed deterioration in separation performance. However, hydrophilic peptides still exhibited poor resolution on both columns. Short ODS-IIIE and SOLAD column backpressures were below 100 bars at 200 nL/min on the Evosep LC (Fig. S4), which is typical for the Whisper methods. Despite the utilization of columns with a diameter twice that of the standard 75 µm dimension, peptide elution was observed within the expected chromatographic range for both preset gradient methods, suggesting that the combination of higher density packing (lower dead volume) and enhanced mass transfer kinetics of nonporous particles effectively compensated for the increased column volume.

**Figure 3.**
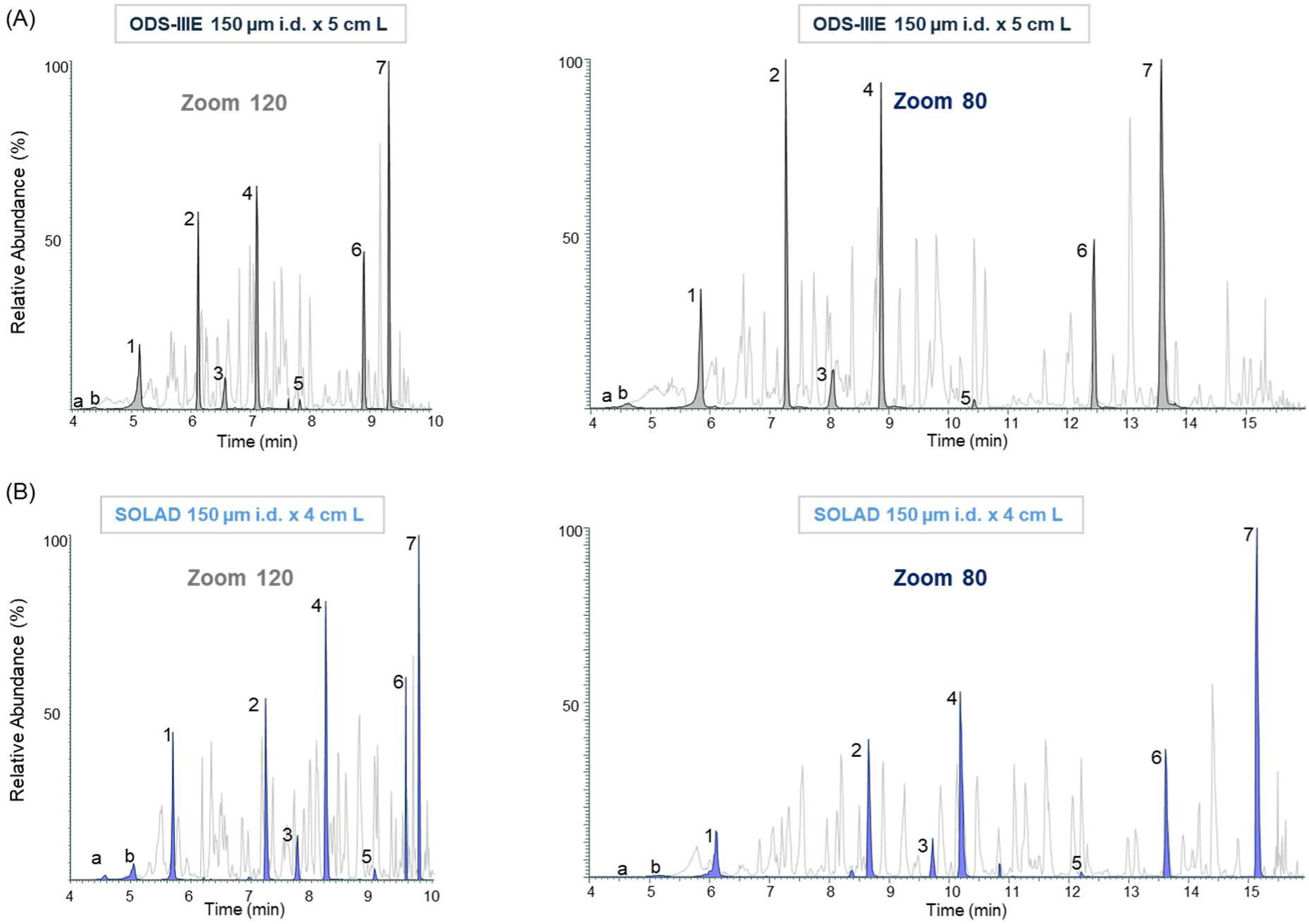
BPCs and XICs of selected tryptic peptides from a 10 fmol bovine serum albumin/α-casein digest separated on a 150 µm i.d. × 5 cm ODS-IIIE column (A) and a 150 µm i.d. × 4 cm SOLAD column (B) using Whisper Zoom 120 (left) and Zoom 80 (right) methods on an Evosep One LC system. Mass spectrometry detection was performed using a Q Exactive instrument.

### Performance Evaluation of short Nonporous and Porous C-18 Columns for Peptide Separation in Fast Liquid-Chromatography

Having established optimal operating conditions for the NPP columns and their feasibility for use with an Evosep LC, we proceeded to compare the separation performance of the short nonporous ODS-IIIE and SOLAD columns with three commonly utilized C-18 materials for peptide separation: a 1.6 µm FPP Luna Omega Polar (75 µm i.d. x 5 cm), a 1.9 µm Reprosil Gold (75 µm i.d. x 5 cm), and a 1.7 µm [1.25 µm (core) and 0.23 µm (shell thickness)] SPP Kinetex (100 µm i.d. x 5 cm), all designed to work at similar pressure range on Evosep system (Fig. 1 and S4). We conducted LC-MS/MS analyses of the same standard digest using the ODS-IIIE, SOLAD, Luna, Reprosil, and Kinetex columns with Whisper Zoom 120 and Zoom 80 methods, and the separation performance of each column was evaluated by assessing the FWHM values of the seven selected peptide peaks (Fig. 4 and Table S2). Visual inspection of the base-peak and extracted ion chromatograms (Fig. 4A) revealed similar elution profiles across all columns under both gradient methods. As expected for the longer gradient, Zoom 80 exhibited broader and more widely spaced peaks. Differences in elution time for the same peptide across different columns, observed in both methods, likely arise from variations in stationary phase chemistries and/or column volumes. At Zoom 120, the overall separation performance was similar across the column materials, with FWHM values ranging from 1.6 to 2.0 seconds (Fig. 4B). The Kinetex column was an exception, exhibiting a slightly higher FWHM of 2.5 seconds. Notably, peptides separated by ODS-IIIE and SOLAD columns exhibited a FWHM of 1.8 and 1.6 seconds, respectively. The longer gradient of Zoom 80, as expected, resulted in higher FWHM values relative to Zoom 120. Under these conditions, ODS-IIIE, SOLAD, and Luna exhibited lower FWHM values (2.9 s, 2.6 s, and 2.6 s, respectively), indicating superior performance compared to Reprosil (3.4 s) and Kinetex (3.8 s) columns. Based on this comparative analysis, the short 150 µm i.d. ODS-IIIE and SOLAD nonporous particle columns demonstrate competitive performance with currently popular materials commonly employed for high throughput peptide separation under equivalent gradient and pressure conditions.

**Figure 4.**
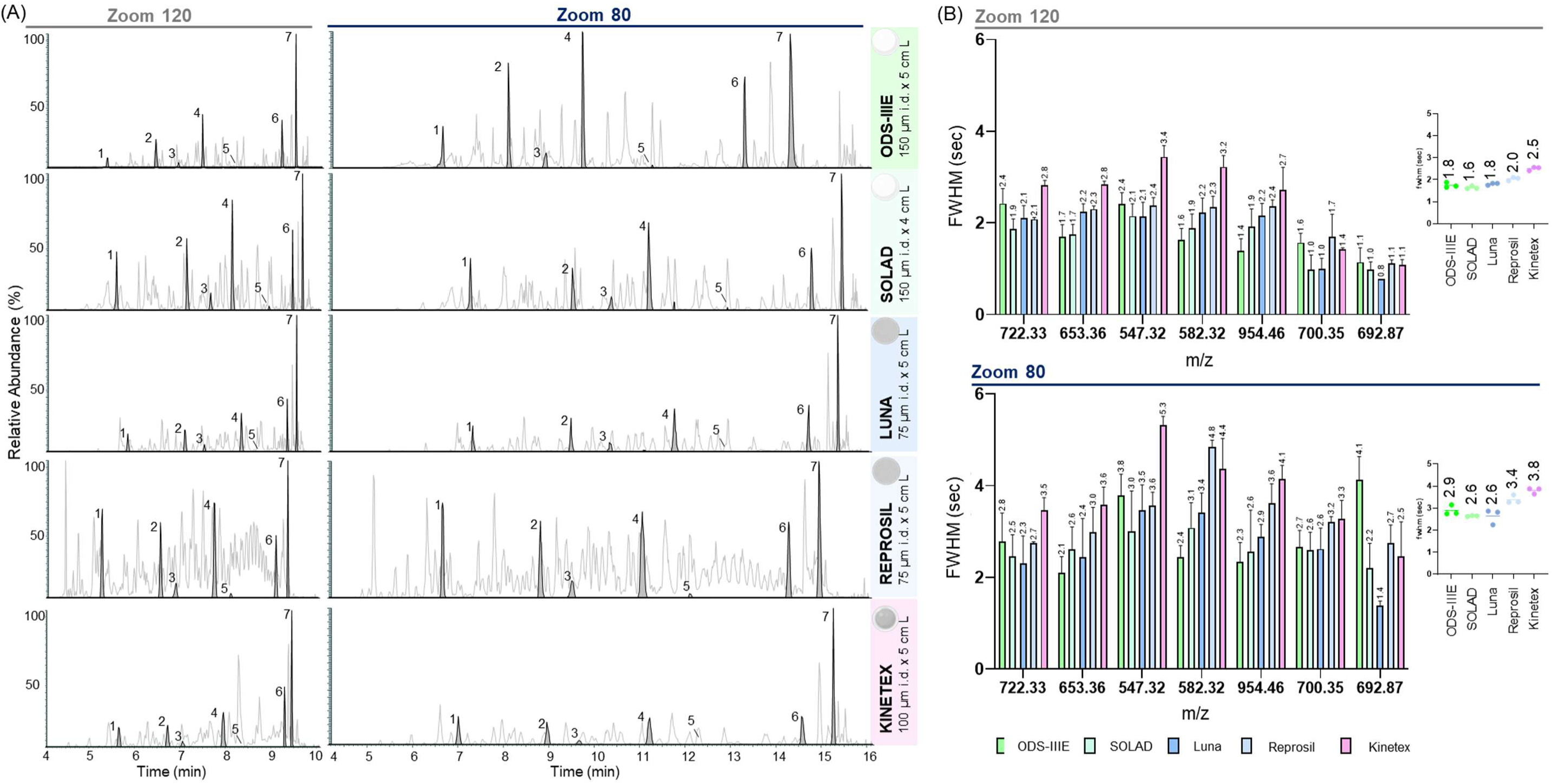
Performance of NPP, FPP, and SPP columns for high-throughput LC-MS/MS using an Evosep One LC system. (A) BPCs and XICs of selected tryptic peptides from a 10 fmol bovine serum albumin/α-casein digest separated using Whisper Zoom 120 (left) and Zoom 80 (right) methods. (B) Individual and average FWHM are presented to indicate the separation efficiency achieved with each column type. Error bars show the standard deviation. Mass spectrometry detection was performed using a Q Exactive.

### Evaluation of Peptide Loading Capacity for Short Nonporous C-18 Columns

While a common concern regarding nonporous materials is their potentially lower binding capacity compared to traditional porous particles^23^, this limitation has become less critical with advancements in mass spectrometer sensitivity.^1, 2^ Nevertheless, we also wanted to evaluate the loading capacity of the ODS-IIIE and SOLAD columns. To this end, we performed LC-MS/MS analyses of increasing amounts (0.25 to 50 ng) of Expi 293F digest with both Whisper Zoom methods and monitored the intensity and FWHM of three representative peptides – an early-, a mid-, and a late-eluting species – as proxies for determining the optimal loading capacity before column performance is compromised by saturation effects. It is well established that mass overloading can lead to peak shape distortion and signal saturation, ultimately compromising data quality and peptide identification.^38, 39^ The loading capacity analysis (Fig. S5 and Table S3) revealed that for both NPP columns and gradient methods, signal intensity for all three monitored peptides began to plateau above approximately 25 ng, a trend consistent with the overall peak intensities observed in the corresponding BPCs shown in Figures S6 and S7. While early- and mid-eluting peptides showed minimal changes in peak width upon saturation, the late-eluting peptide exhibited a marked increase in FWHM when the injected mass exceeded 25 ng, consistent with the observation that the overloading effects intensify with the increase in the retention factor *k*.^40^ Based on these trends, a loading amount below 25 ng of the human cell lysate digest appears to be optimal for both ODS-IIIE and SOLAD columns under both gradient methods to preserve peak shape and prevent significant intensity saturation. Such amounts are compatible/optimal with most modern mass spectrometers.

### Performance of Nonporous and Porous Columns in High-Throughput DIA LC-MS/MS

Having established the peptide loading capacity of short ODS-IIIE and SOLAD columns, we proceeded to evaluate their performance against the other commonly utilized columns (Luna, Reprosil and Kinetex) in a high-throughput DIA workflow using the Whisper Zoom 120 and 80 methods. To ensure a fair comparison, we selected a loading amount of 20 ng of Expi 293F digest, below the established saturation threshold observed for the short NPP columns. Using the Zoom 120 method, precursor identification was relatively consistent across the five columns, ranging from 19,740 (Kinetex) to 23,100 (SOLAD), and showed good reproducibility across technical replicates, as evidenced by the relatively small variations (Fig. 5 and Table S4). Protein identification was also comparable and ranged from 2,707 to 3,130. Notably, nonporous particle columns (ODS-IIIE and SOLAD) yielded the highest numbers of proteins (>3,000), followed by the fully porous particle columns, Luna (2,942) and Reprosil (2,977). The lowest protein count was observed for the superficially porous Kinetex column (2,707). As expected, the longer gradient provided by the Zoom 80 method, significantly increased both precursor and protein identification, with average improvements of 40% and 20%, respectively, relative to Zoom 120. Precursor counts with Zoom 80 ranged from 27,353 (Kinetex) to 32,813 (SOLAD), while protein counts ranged from 3,327 (Kinetex) to 3,881 (ODS-IIIE). The same trend in protein identification performance observed with the Zoom 120 method was also seen with the Zoom 80 method: ODS-IIIE and SOLAD columns yielded the highest protein counts (>3,800). Based on these observations, both precursor and protein identification were consistent and reproducible for all columns tested across both gradient methods. The nonporous ODS-IIIE and SOLAD columns demonstrated comparable or superior performance to the fully and superficially porous columns using high-throughput DIA workflow.

**Figure 5.**
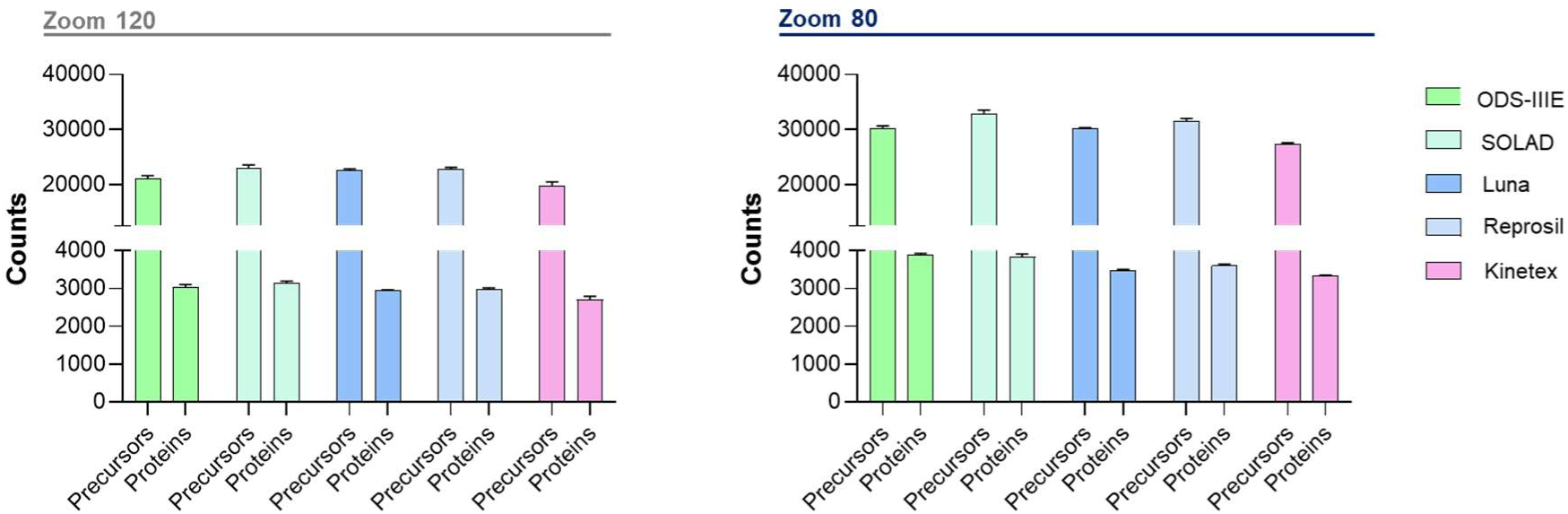
Evaluation of peptide and protein identification performance for the analysis of 20 ng of Expi 293F digest using the Evosep One LC system with Zoom 120 and Zoom 80 methods, coupled with DIA on an Orbitrap Exploris 480 mass spectrometer. The average number is represented by the bars, and error bars show the standard deviation calculated from three independent LC-MS/MS analyses.

## Conclusions

We demonstrate that short nonporous C-18 columns packed with sub-2 µm particles offer competitive chromatographic performance for high-speed nanoLC-based proteomics workflows. We show that the long and short versions of these columns enable sharp peak profiles and efficient peptide separations within pressure limits compatible with widely used nanoLC systems. Comparative analyses reveal that nonporous columns deliver similar or superior separation performance relative to established fully and superficially porous C-18 materials, while maintaining competitive protein/precursor identifications under high-throughput DIA conditions using 20 ng of cell lysate digest. These findings highlight the potential of nonporous C-18 phases as effective alternatives for rapid, sensitive proteomic analyses using short gradients and low sample input conditions.

## AUTHOR INFORMATION

### Author contributions

E.S.K. and S.M.: Conceptualization; E.S.K. and Y.D. performed the experiments; E.S.K. collected the data; E.S.K. and S.M. analysed the data; E.S.K. and S.M. wrote the initial draft; All authors read, edited and approved the final version of the paper.

### Notes

The authors declare no competing financial interest(s).

## ACKNOWLEDGMENTS

S.M, E.S.K and Y.D, were supported by EPSRC (V011359/1 (P)).

## Supporting Information

### METHOD

#### Protein Digestion and Sample Preparation

Bovine serum albumin (BSA, Sigma) was solubilized in 1 M urea, 100 mM ammonium bicarbonate (AmBic), while α-casein (Sigma) was dissolved in 8 M urea, 100 mM AmBic. BSA and casein samples were reduced with 10 mM tris(2-carboxyethyl)phosphine (TCEP) for 5 minutes at room temperature and alkylated with 50 mM 2-chloroacetamide (CAA, Sigma) for 30 minutes in the dark. The casein sample was then diluted with 100 mM AmBic to a final urea concentration of 4 M. Both BSA and casein samples were subsequently digested with Lys-C (Wako, biochemistry grade) with a protein-to-enzyme ratio of 100:1 (w/w) for 2 hours at 37 °C. Following Lys-C digestion, the samples were further diluted with 100 mM AmBic to achieve final urea concentration of approximately 1 M for casein and 0.25 M for BSA. Trypsin digestion (Promega) was performed at 37 °C using enzyme-to-substrate ratios of 1:30 (casein) and 1:40 (BSA), with an incubation time of 4 hours. After digestion, samples were acidified with formic acid (FA) and cleaned up using Oasis® HLB cartridges (3 cc, 60 mg sorbent, Waters). Eluted peptides were dried using a Genevac concentrator. Casein and BSA digests were mixed and diluted in either 5% FA / 5% dimethyl sulfoxide (DMSO) to a final concentration of 10 fmol/µL for nanoUPLC analyses, or with 5% FA / 0.015% n-Dodecyl-β-D-maltoside (DDM) to a final concentration of 2 fmol/µL prior to loading onto Evotip pure (Evosep Biosystems) for subsequent nanoHPLC analysis.

Expi293F cells (Gibco) were resuspended in a lysis buffer composed of 8 M urea, 100 mM AmBic, and a protease inhibitor cocktail (Roche). Samples were sonicated at 4 °C using a Bioruptor Ultrasonicator (Bioruptor Pico, Diagenode) set to ultra-low frequency for 30 cycles of 30 seconds each. After centrifugation at 14,000 x g, 4 °C for 15 minutes, the supernatants were collected, and the protein concentrations were estimated by BCA assay (Pierce). Proteins were reduced with 10 mM TCEP for 30 minutes at room temperature followed by alkylation with 50 mM CAA for 30 minutes in the dark. The urea concentration was diluted to 4 M with 100 mM AmBic, and the first step of digestion was carried out with Lys-C protease for 2 hours at 37 °C with a protein-to-enzyme ratio of 50:1 (w/w). Samples were further diluted to 1.3 M urea and digested with trypsin for 4 hours at 37 °C with a protein-to-enzyme ratio of 50:1 (w/w). Peptide samples were cooled to 4 °C, acidified with FA to a final concentration of 5% (v/v) and centrifuged at 16,200 × g for 10 minutes to remove any precipitate. The supernatants were diluted in 5% FA / 0.015% DDM at concentrations ranging from 0.0125 to 2.5 ng/µL prior to loading onto Evotip pure for subsequent nanoHPLC analysis.

Peptide samples analysed using the Evosep One HPLC system (Evosep Biosystems) were prepared with Evotips following the manufacturer’s instructions.

#### Column Packing

Nonporous (NPP), fully porous (FPP), and superficially porous C-18 particles (SPP) were prepared as slurries at a concentration of 100 mg/mL. FPP Reprosil Gold™, and the NPPs ODS-IIIE™ and SOLAD™ were slurried in 100% acetone, while Luna Omega Polar™ (FPP) and Kinetex® (SPP) were slurried in 100% methanol. Prior to packing, the materials were washed three times with either acetone or methanol, followed by 20 minutes of sonication and particle sedimentation. Fused silica capillaries (Polymicro Technologies) with internal diameters (i.d.) of 75, 100 and 150 µm, and outer diameter (o.d.) of 360 µm were pulled using a P-2000 laser puller (Sutter Instrument). Laser parameters were set as follows: HEAT = 245, FIL = -, VEL = 15, DEL = 132, PUL = - for the 75 µm i.d. capillaries, and HEAT = 300, FIL = -, VEL = 15, DEL = 255, PUL = 10 for the 100 and 150 µm i.d. capillaries. Capillaries were then packed with the respective C-18 materials using a PC8500 Pressure Injection Cell (Next Advance), following the protocol described by Kovalchuck^1^, with the production of a self-assembled particles as a frit at the outlet of the columns.^2^ Seventy-five micrometre i.d. capillaries were packed with fully porous Reprosil Gold™ (1.9 µm, Dr. Maisch) and Luna Omega Polar™ (1.6 µm, Phenomenex); 100 µm i.d. capillary was packed with superficially porous Kinetex® [1.7 µm – 1.25 µm (core) and 0.23 µm (shell thickness) – Phenomenex)]; and 150 µm i.d. capillaries, with nonporous ODS-IIIE™ (1.5 µm, Eprogen/Promigen Life Sciences) and SOLAD™ (1.0 µm, Glantreo). Columns were packed at 120 bar for a minimum of 2 hours, after which the pressure was gently released. To consolidate the column beds, a constant flow of 70% acetonitrile (ACN) was applied at 800 bar for 2 hours using an NCS-3500RS nano pump (Thermo Scientific). Following consolidation, columns were cut to the desired lengths: 25 cm and 5 cm for ODS-IIIE; 15 cm and 4 cm for SOLAD; and 5 cm for Luna, Reprosil, and Kinetex. A sol-gel frit was formed at the column inlet according to the method described by Maiolica.^3^ Briefly, the capillary inlets were gently pushed on a glass microfibre filter (GF/C, Whatman) previously wet with a 1:1 (v/v) mixture of Kasil 1624 (Next Advance) and 25% formamide. The frits were then polymerized at 85 °C for 16 hours. LC-MS/MS analyses were performed using three independent 5-cm ODS-IIIE and 4-cm SOLAD columns, while experiments involving the longer NPP columns and the 5-cm Luna, Reprosil, and Kinetex columns, were carried out using a single column of each type.

#### Nano-Liquid Chromatography and Mass Spectrometry

To optimize flow rates and column temperatures for the NPP C-18 columns, 10 fmol BSA/casein digest was analysed using an Ultimate 3000 RSLCnano system (Thermo Fisher Scientific) equipped with a C-18 PepMap100 trap column (300 μm i.d. x 5 mm L, 100Å, Thermo Fisher Scientific), and separated on 25-cm ODS-IIIE and 15-cm SOLAD analytical columns. Solvent A was composed of 5% DMSO and 0.1% formic acid in water, while solvent B contained 5% DMSO and 0.1% FA in ACN. Chromatographic separations were conducted over a temperature range of 30 – 70 °C at a constant flow rate of 200 nL/min, and at flow rates ranging from 50 to 500 nL/min at 40 °C (ODS-IIIE) and 50 °C (SOLAD), utilizing a 15-minute linear gradient as follows: a 5-minute hold at 2% B, and 7% to 55% solvent B over 15 minutes, followed by an increase to 99% B over 2 minutes, a 1-minute hold at 99% B, and re-equilibration to 2% B for variable duration depending on the employed flow rate. The BSA/casein digest was also subjected to a 15-minute separation using shorter ODSIII (5 cm) and SOLAD (4 cm) columns at 200 nL/min at 20 °C. To assess and compare the separation performance of ODS-IIIE and SOLAD columns with that of FPP (Luna and Reprosil) and SPP (Kinetex) columns, a 10 fmol BSA/casein digest was analysed using the predefined Whisper Zoom 120 (10-minute gradient) and Zoom 80 (16-minute gradient) methods on an Evosep One LC. Increasing amounts of Expi 293F digest (0.25 to 50 ng) separated by the Whisper Zoom 120 and Zoom 80 methods was used for the assessment of peptide loading capacity of 5-cm ODSIII and 4-cm SOLAD columns. Data-Dependent Acquisition (DDA) mass spectrometry was performed on a Q Exactive mass spectrometer (Thermo Fisher Scientific) using the following parameters: Full MS scans (350-1,400 m/z) were acquired in the Orbitrap at 70,000 resolution (at m/z 200) using an AGC target of 3 × 10⁶ and a maximum injection time of 50 ms. The top five most intense precursors (charge states 2–7) were isolated with a 1.5 Th window and fragmented using higher-energy collisional dissociation (HCD) at a normalized collision energy (NCE) of 30%. MS/MS spectra were acquired in the Orbitrap at 17,500 resolution with an AGC target of 5 × 10⁴ and a maximum injection time of 128 ms. Dynamic exclusion was enabled with a 5 s exclusion duration, ±10 ppm mass tolerance, and a repeat count of 1.

Twenty nanograms of Expi 293F digest were separated on 5-cm NPP (150 μm i.d.), FPP (75 μm i.d.), and SPP (100 μm i.d.) columns, except for the SOLAD material, where a 4 cm column was used. Separations were carried out using the Whisper Zoom 120 and Zoom 80 methods on an Evosep One system coupled to an Orbitrap Exploris 480 mass spectrometer (Thermo Fisher Scientific) operating in DIA mode. Full MS scans (350−1400 m/z) were acquired at 60,000 resolution with a 300% normalised AGC target, and 25 ms injection time. DIA was performed using 33 isolation windows (12 m/z width) spanning the m/z range of 400–800. Precursor ions were fragmented using HCD with a NCE of 30%. MS/MS spectra were acquired in the Orbitrap at 15,000 resolution with a normalized AGC target of 2000% and injection time of 22 ms. The total cycle time was 3 seconds.

LC-MS/MS analyses were conducted in triplicate for each column, except for the column temperature evaluation, where a single analysis was performed per column at each tested temperature.

#### Raw Data Analysis

Full-width at half maximum (FWHM) and peptide intensities were manually calculated/obtained from the extracted ion chromatograms (XICs) of selected peptides from BSA/casein or Expi 293F digests using Freestyle™ software version 1.8.51.0 (Thermo Fisher Scientific). Data were further processed using GraphpadPrism version 10.4.2.

DIA LC-MS/MS data were analysed by DIA-NN software^4^ (version 1.8.1) using library-free search. Spectral library was predicted in silico from the Uniprot human proteome database (Proteome ID: UP000005640, downloaded in August 2022, 79,759 sequences). Methionine oxidation and cysteine carbamidomethylation were set as variable and fixed modification, respectively. Enzyme specificity was set to trypsin/P, allowing up to one missed cleavage, and peptide lengths were restricted to 7–30 amino acids. The following parameters were used: protein inference set to ‘Genes’, neural network classifier set to ‘Single-pass mode’, and quantification strategy selected as ‘Robust LC (high precision)’. Cross-run normalization was performed using the ‘RT-dependent’ setting, and spectral library generation was enabled using the ‘Smart profiling’ mode. Speed and RAM usage were configured for ‘Optimal results’. Mass accuracy settings for both precursor and fragment ions were set to 0 to enable automatic inference. The options ‘No shared spectra’, ‘Heuristic protein inference’, and ‘Unrelated runs’ were enabled. Precursor-level false discovery rate (FDR) was estimated and filtered to 1%. Raw data and search results have been deposited in the ProteomeXchange Consortium via the PRIDE^5^ partner repository under the dataset identifier PXD064019.

### Supplemental Figures

**Figure S1.**
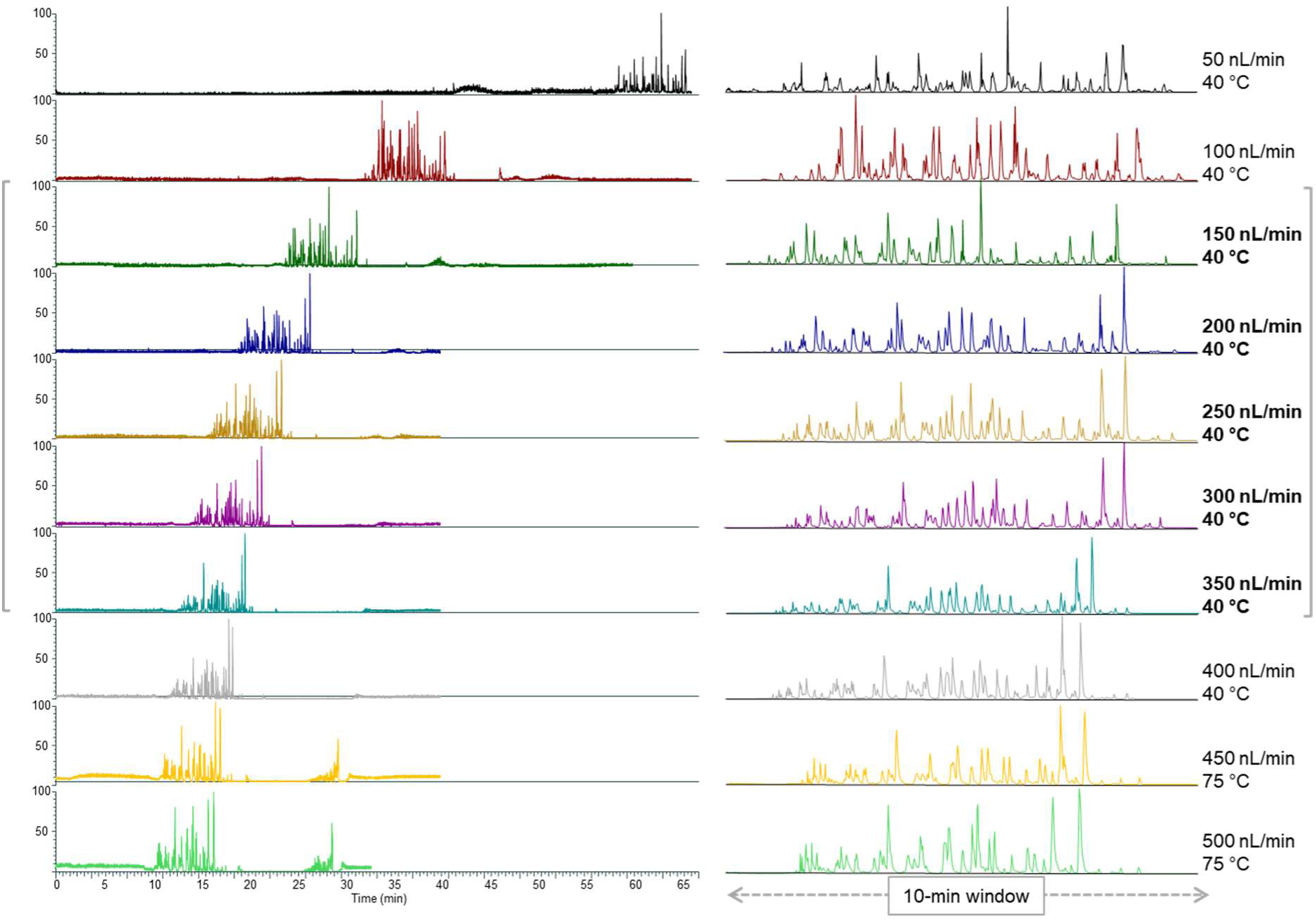
BPCs of a 10 fmol tryptic digest of bovine serum albumin/α-casein separated at increasing flow rates using an Ultimate 3000 RSLCnano system with a 150 µm i.d. × 25 cm ODS-IIIE column and a 15-minute linear gradient (7-55% solvent B). The column temperature was maintained at 40 °C during LC-MS/MS analysis and increased to 75 °C when the flow rate exceeded 400 nL/min to control backpressure. Optimal flow rates (150-350 nL/min) are highlighted in brackets. Detection was performed with a Q Exactive mass spectrometer.

**Figure S2.**
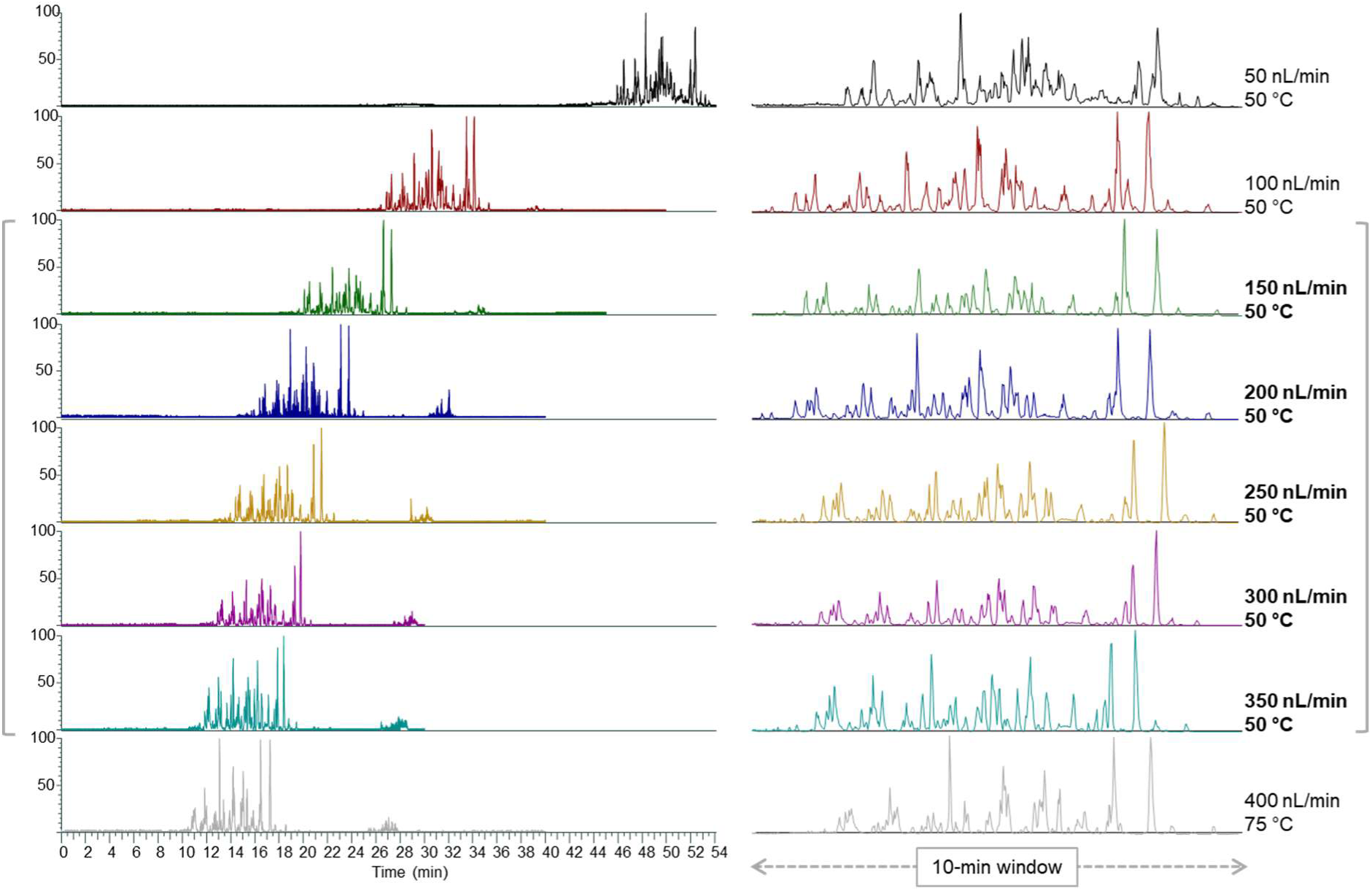
BPCs of a 10 fmol tryptic digest of bovine serum albumin/α-casein separated at increasing flow rates using an Ultimate 3000 RSLCnano system with a 150 µm i.d. × 15 cm SOLAD column and a 15-minute linear gradient (7-55% solvent B). The column temperature was maintained at 50 °C during LC-MS/MS analysis and increased to 75 °C when the flow rate exceeded 350 nL/min to control backpressure. Optimal flow rates (150-350 nL/min) are highlighted in brackets. Detection was performed with a Q Exactive mass spectrometer.

**Figure S3.**
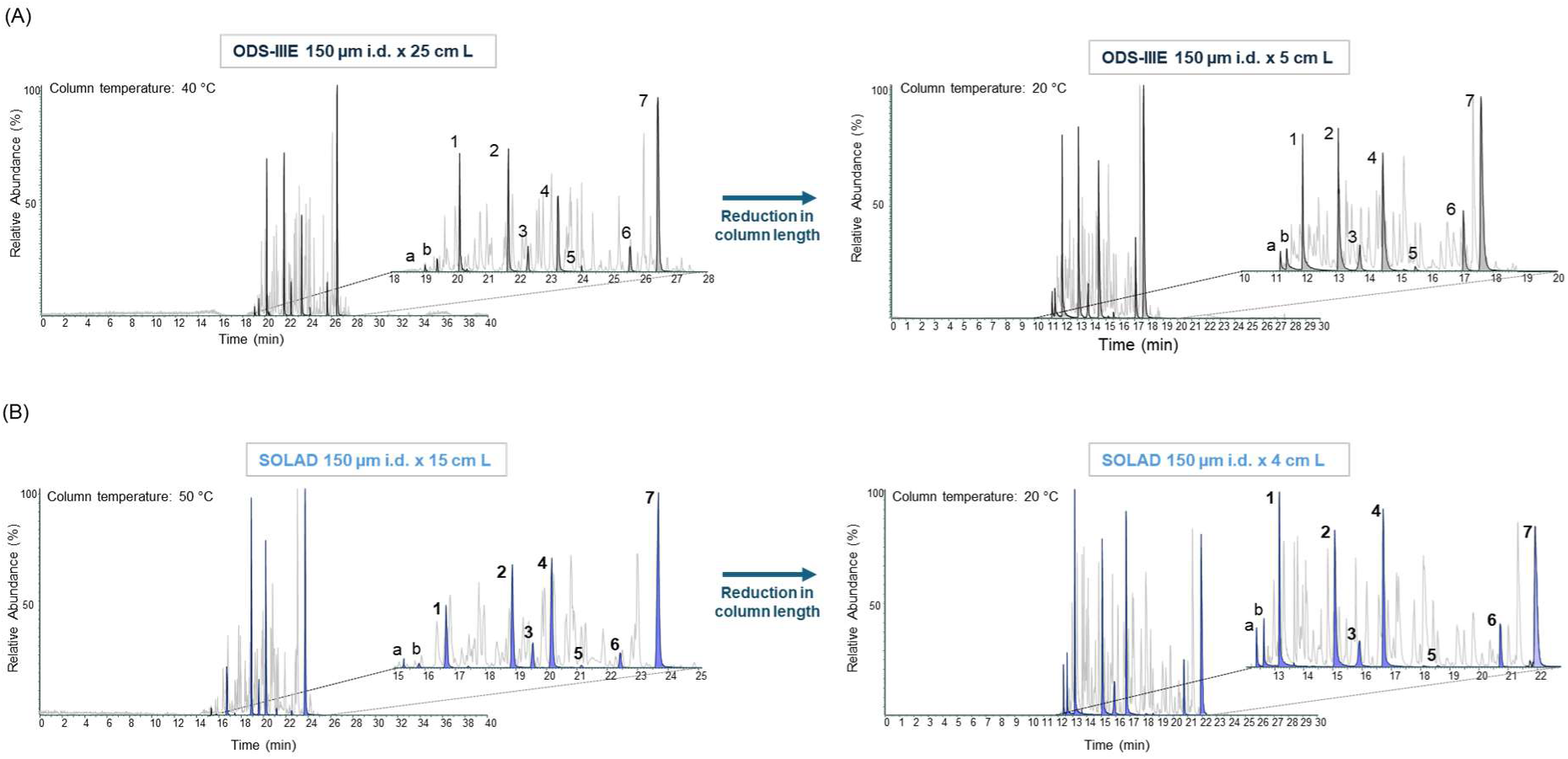
BPCs and XICs of selected peptides from a 10 fmol bovine serum albumin/α-casein tryptic digest, analysed by 15-min gradient LC-MS/MS at a flow rate of 200 nL/min using long and short ODS-IIIE (A) and SOLAD (B) columns. LC-MS/MS was performed on an Ultimate 3000 RSLCnano system coupled to a Q Exactive mass spectrometer. Early-eluting peptides (m/z 488.54 and 625.78) are represented by peaks “a” and “b”, respectively.

**Figure S4.**
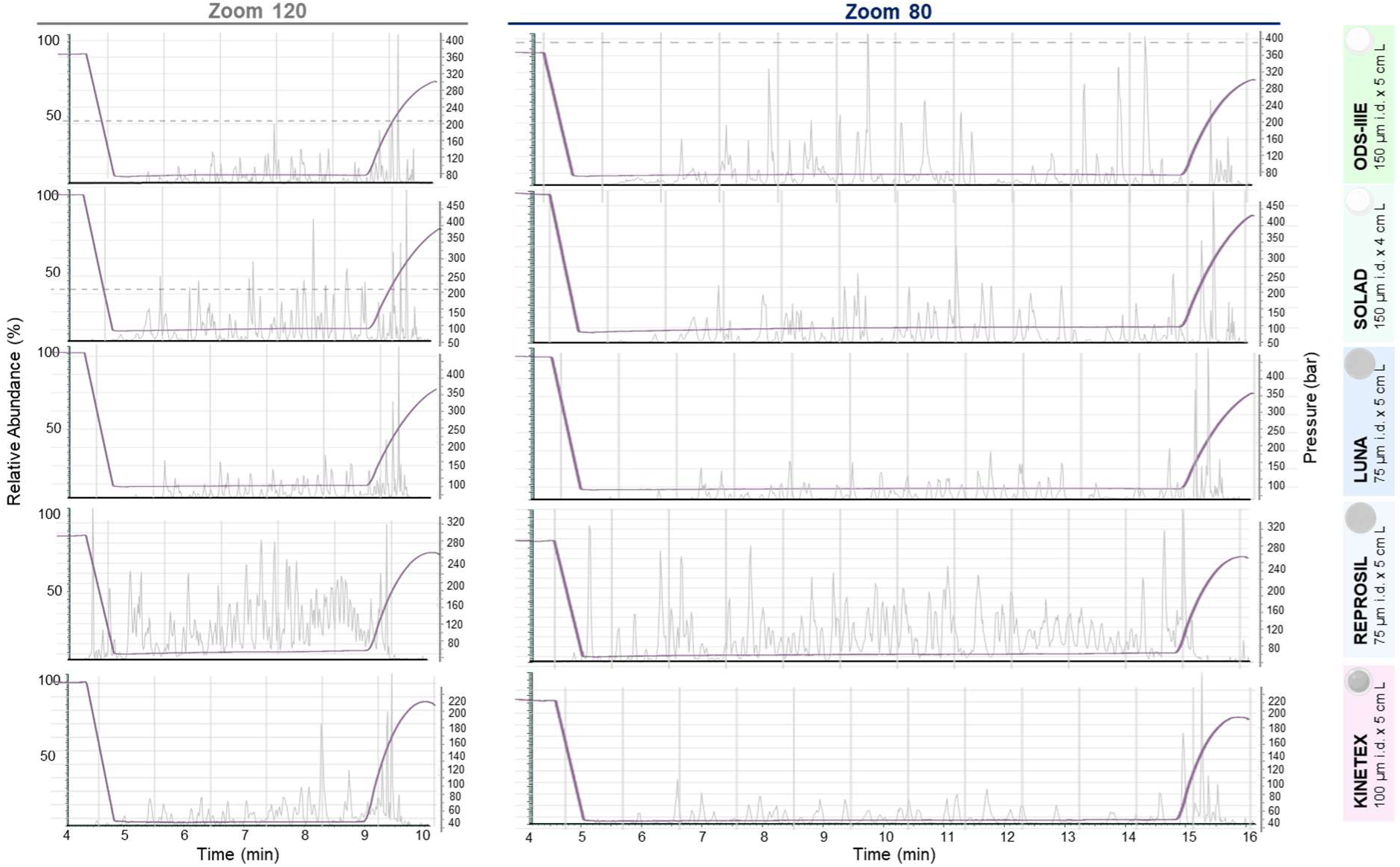
BPCs and high-pressure pump profiles from LC-MS/MS analysis of 10 fmol tryptic peptides derived from a bovine serum albumin/α-casein digest, separated using the Whisper Zoom 120 (left) and Zoom 80 (right) methods with NPP, FPP, and SPP columns.

**Figure S5.**
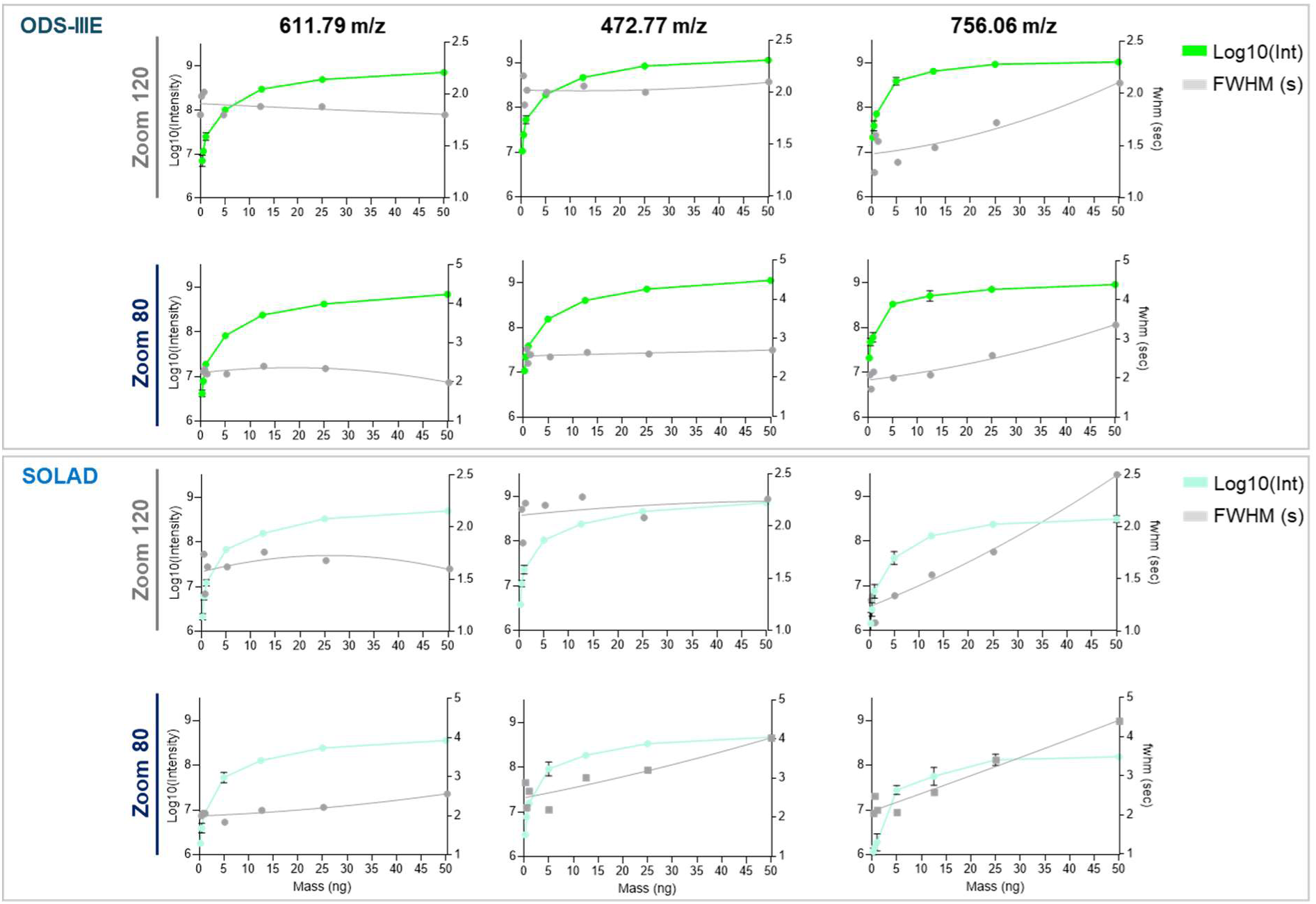
Peptide loading capacity evaluation of 150 µm i.d. × 5 cm L ODS-IIIE (top panel) and 150 µm i.d. × 4 cm L SOLAD (bottom panel) columns. Average intensities and FWHM (n=3) of three peptides are plotted against increasing amounts of Expi 293F digest (0.25 to 50 ng) separated by Whisper Zoom 120 and Zoom 80 methods on an Evosep One LC system. Grey lines indicate second-degree polynomial fits applied to the data points to visualize trends in FWHM. Error bars correspond to the standard deviation.

**Figure S6.**
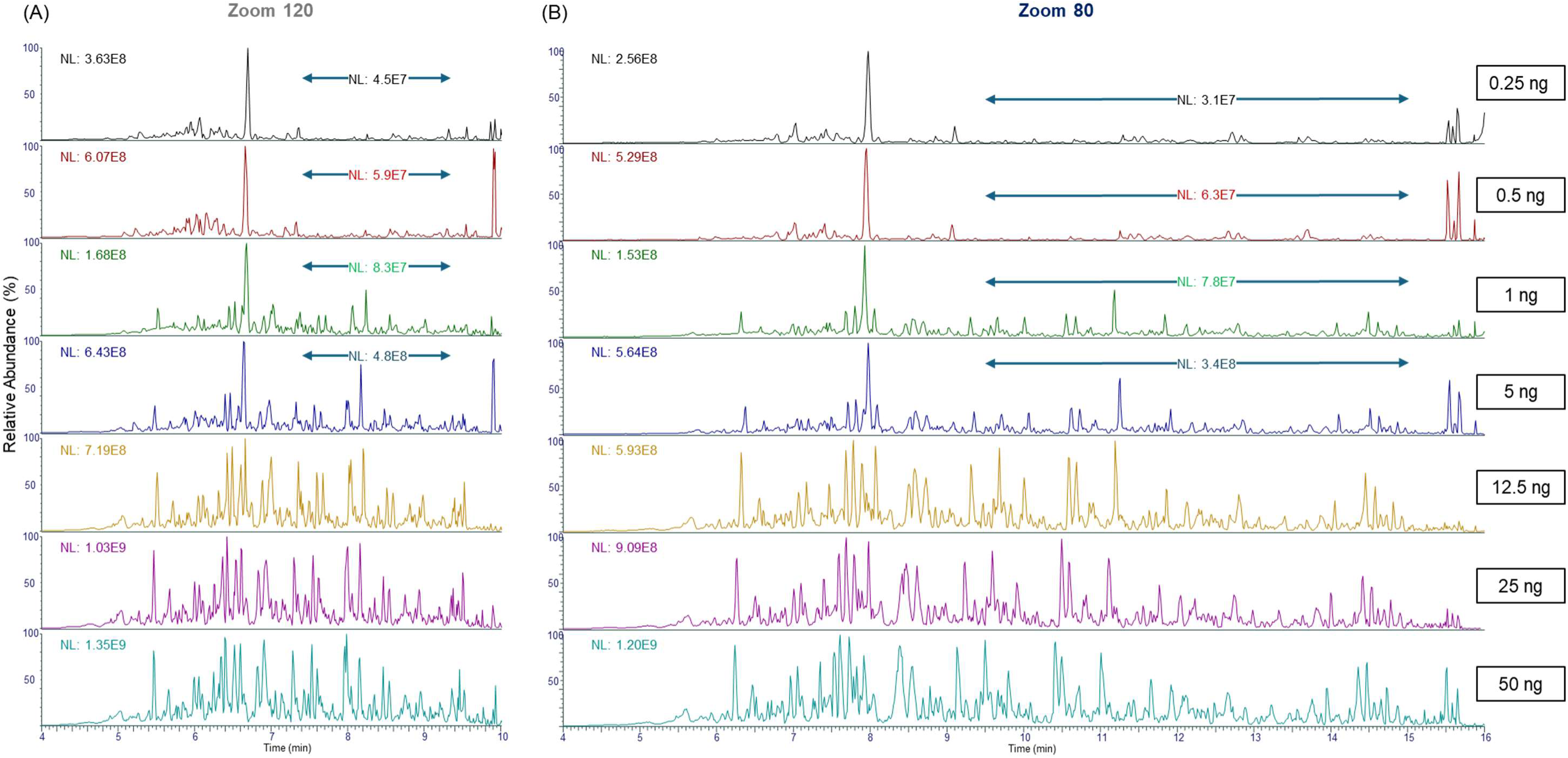
Representative base peak chromatograms obtained from LC-MS/MS analysis of increasing amounts of Expi 293F digest (0.25 – 50 ng) using the Whisper Zoom 120 (A) and Zoom 80 (B) gradient methods on an Evosep LC system equipped with a 150 µm i.d. × 5 cm L ODS-IIIE column. Double-arrow lines indicate regions that are magnified to provide a clearer visualization of the average peptide intensities. Mass spectrometry detection was performed using a Q Exactive system.

**Figure S7.**
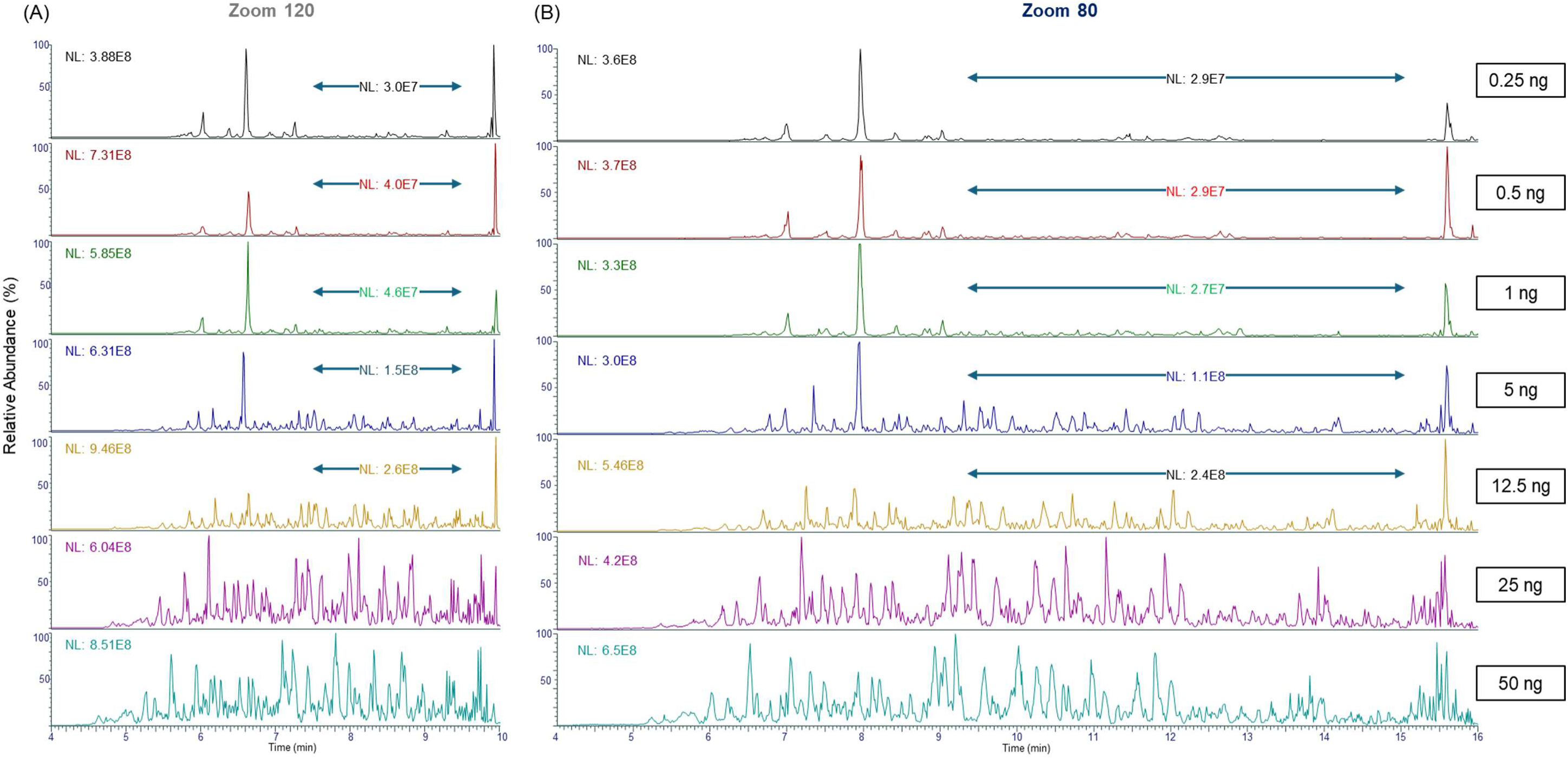
Representative base peak chromatograms obtained from LC-MS/MS analysis of increasing amounts of Expi 293F digest (0.25 – 50 ng) using the Whisper Zoom 120 (A) and Zoom 80 (B) gradient methods on an Evosep LC system equipped with a 150 µm i.d. × 4 cm L SOLAD column. Double-arrow lines indicate regions that are magnified to provide a clearer visualization of the average peptide intensities. Mass spectrometry detection was performed using a Q Exactive system.

## References

(1) Meier, F.; Brunner, A. D.; Koch, S.; Koch, H.; Lubeck, M.; Krause, M.; Goedecke, N.; Decker, J.; Kosinski, T.; Park, M. A.;, et al. Online Parallel Accumulation-Serial Fragmentation (PASEF) with a Novel Trapped Ion Mobility Mass Spectrometer. Mol Cell Proteomics 2018, 17 (12), 2534–2545. DOI: 10.1074/mcp.TIR118.000900 From NLM.

(2) Heil, L. R.; Damoc, E.; Arrey, T. N.; Pashkova, A.; Denisov, E.; Petzoldt, J.; Peterson, A. C.; Hsu, C.; Searle, B. C.; Shulman, N.;, et al. Evaluating the Performance of the Astral Mass Analyzer for Quantitative Proteomics Using Data-Independent Acquisition. J Proteome Res 2023, 22 (10), 3290–3300. DOI: 10.1021/acs.jproteome.3c00357 From NLM.

(3) Bache, N.; Geyer, P. E.; Bekker-Jensen, D. B.; Hoerning, O.; Falkenby, L.; Treit, P. V.; Doll, S.; Paron, I.; Müller, J. B.; Meier, F.;, et al. A Novel LC System Embeds Analytes in Pre-formed Gradients for Rapid, Ultra-robust Proteomics. Mol Cell Proteomics 2018, 17 (11), 2284–2296. DOI: 10.1074/mcp.TIR118.000853 From NLM.

(4) Kreimer, S.; Haghani, A.; Binek, A.; Hauspurg, A.; Seyedmohammad, S.; Rivas, A.; Momenzadeh, A.; Meyer, J. G.; Raedschelders, K.; Van Eyk, J. E. Parallelization with Dual-Trap Single-Column Configuration Maximizes Throughput of Proteomic Analysis. Analytical Chemistry 2022, 94 (36), 12452–12460. DOI: 10.1021/acs.analchem.2c02609.

(5) Webber, K. G. I.; Truong, T.; Johnston, S. M.; Zapata, S. E.; Liang, Y.; Davis, J. M.; Buttars, A. D.; Smith, F. B.; Jones, H. E.; Mahoney, A. C.;, et al. Label-Free Profiling of up to 200 Single-Cell Proteomes per Day Using a Dual-Column Nanoflow Liquid Chromatography Platform. Analytical Chemistry 2022, 94 (15), 6017–6025. DOI: 10.1021/acs.analchem.2c00646.

(6) Xie, X.; Truong, T.; Huang, S.; Johnston, S. M.; Hovanski, S.; Robinson, A.; Webber, K. G. I.; Lin, H.-J. L.; Mun, D.-G.; Pandey, A.;, et al. Multicolumn Nanoflow Liquid Chromatography with Accelerated Offline Gradient Generation for Robust and Sensitive Single-Cell Proteome Profiling. Analytical Chemistry 2024, 96 (26), 10534–10542. DOI: 10.1021/acs.analchem.4c00878.

(7) Kverneland, A. H.; Harking, F.; Vej-Nielsen, J. M.; Huusfeldt, M.; Bekker-Jensen, D. B.; Svane, I. M.; Bache, N.; Olsen, J. V. Fully Automated Workflow for Integrated Sample Digestion and Evotip Loading Enabling High-Throughput Clinical Proteomics. Molecular & Cellular Proteomics 2024, 23 (7). DOI: 10.1016/j.mcpro.2024.100790 (acccessed 2025/04/22).

(8) Hansen, C. B.; Møller, M. E. E.; Pérez-Alós, L.; Israelsen, S. B.; Drici, L.; Ottenheijm, M. E.; Nielsen, A. B.; Wewer Albrechtsen, N. J.; Benfield, T.; Garred, P. Differences in biomarker levels and proteomic survival prediction across two COVID-19 cohorts with distinct treatments. iScience 2025, 28 (3), 112046. DOI: 10.1016/j.isci.2025.112046.

(9) Messner, C. B.; Demichev, V.; Bloomfield, N.; Yu, J. S. L.; White, M.; Kreidl, M.; Egger, A.-S.; Freiwald, A.; Ivosev, G.; Wasim, F.;, et al. Ultra-fast proteomics with Scanning SWATH. Nature Biotechnology 2021, 39 (7), 846–854. DOI: 10.1038/s41587-021-00860-4.

(10) Niu, L.; Thiele, M.; Geyer, P. E.; Rasmussen, D. N.; Webel, H. E.; Santos, A.; Gupta, R.; Meier, F.; Strauss, M.; Kjaergaard, M.;, et al. Noninvasive proteomic biomarkers for alcohol-related liver disease. Nat Med 2022, 28 (6), 1277–1287. DOI: 10.1038/s41591-022-01850-y From NLM.

(11) Ye, Z.; Sabatier, P.; van der Hoeven, L.; Lechner, M. Y.; Phlairaharn, T.; Guzman, U. H.; Liu, Z.; Huang, H.; Huang, M.; Li, X.;, et al. Enhanced sensitivity and scalability with a Chip-Tip workflow enables deep single-cell proteomics. Nature Methods 2025. DOI: 10.1038/s41592-024-02558-2.

(12) Ctortecka, C.; Clark, N. M.; Boyle, B. W.; Seth, A.; Mani, D. R.; Udeshi, N. D.; Carr, S. A. Automated single-cell proteomics providing sufficient proteome depth to study complex biology beyond cell type classifications. Nature Communications 2024, 15 (1), 5707. DOI: 10.1038/s41467-024-49651-w.

(13) Ai, L.; Binek, A.; Zhemkov, V.; Cho, J. H.; Haghani, A.; Kreimer, S.; Israely, E.; Arzt, M.; Chazarin, B.; Sundararaman, N.;, et al. Single Cell Proteomics Reveals Specific Cellular Subtypes in Cardiomyocytes Derived from Human iPSCs and Adult Hearts. Molecular & Cellular Proteomics 2025, 100910. DOI: 10.1016/j.mcpro.2025.100910.

(14) Gritti, F.; Guiochon, G. The current revolution in column technology: How it began, where is it going? Journal of Chromatography A 2012, 1228, 2–19. DOI: 10.1016/j.chroma.2011.07.014.

(15) Shishkova, E.; Hebert, A. S.; Coon, J. J. Now, More Than Ever, Proteomics Needs Better Chromatography. Cell Syst 2016, 3 (4), 321–324. DOI: 10.1016/j.cels.2016.10.007 From NLM.

(16) Hayes, R.; Ahmed, A.; Edge, T.; Zhang, H. Core–shell particles: Preparation, fundamentals and applications in high performance liquid chromatography. Journal of Chromatography A 2014, 1357, 36–52. DOI: 10.1016/j.chroma.2014.05.010.

(17) Koshiyama, A.; Ichibangase, T.; Moriya, K.; Koike, K.; Yazawa, I.; Imai, K. Liquid chromatographic separation of proteins derivatized with a fluorogenic reagent at cysteinyl residues on a non-porous column for differential proteomics analysis. Journal of Chromatography A 2011, 1218 (22), 3447–3452. DOI: 10.1016/j.chroma.2011.03.070.

(18) Kawashima, Y.; Ohara, O. Development of a NanoLC–MS/MS System Using a Nonporous Reverse Phase Column for Ultrasensitive Proteome Analysis. Analytical Chemistry 2018, 90 (21), 12334–12338. DOI: 10.1021/acs.analchem.8b03382.

(19) Schuster, S. A.; Boyes, B. E.; Wagner, B. M.; Kirkland, J. J. Fast high performance liquid chromatography separations for proteomic applications using Fused-Core® silica particles. Journal of Chromatography A 2012, 1228, 232–241. DOI: 10.1016/j.chroma.2011.07.082.

(20) Issaeva, T.; Kourganov, A.; Unger, K. Super-high-speed liquid chromatography of proteins and peptides on non-porous Micra NPS-RP packings. Journal of Chromatography A 1999, 846 (1), 13–23. DOI: 10.1016/S0021-9673(99)00360-X.

(21) Wu, N.; Liu, Y.; Lee, M. L. Sub-2μm porous and nonporous particles for fast separation in reversed-phase high performance liquid chromatography. Journal of Chromatography A 2006, 1131 (1), 142–150. DOI: 10.1016/j.chroma.2006.07.042.

(22) Fekete, S.; Guillarme, D. Possibilities of new generation columns packed with 1.3μm core–shell particles in gradient elution mode. Journal of Chromatography A 2013, 1320, 86–95. DOI: 10.1016/j.chroma.2013.10.061.

(23) Jorgenson, J. W. Capillary liquid chromatography at ultrahigh pressures. Annu Rev Anal Chem (Palo Alto Calif) 2010, 3, 129–150. DOI: 10.1146/annurev.anchem.1.031207.113014 From NLM.

(24) Kirkland, J. J.; Schuster, S. A.; Johnson, W. L.; Boyes, B. E. Fused-core particle technology in high-performance liquid chromatography: An overview. J Pharm Anal 2013, 3 (5), 303–312. DOI: 10.1016/j.jpha.2013.02.005 From NLM.

(25) Lenčo, J.; Jadeja, S.; Naplekov, D. K.; Krokhin, O. V.; Khalikova, M. A.; Chocholouš, P.; Urban, J.; Broeckhoven, K.; Nováková, L.; Švec, F. Reversed-Phase Liquid Chromatography of Peptides for Bottom-Up Proteomics: A Tutorial. J Proteome Res 2022, 21 (12), 2846–2892. DOI: 10.1021/acs.jproteome.2c00407 From NLM.

(26) Fröhlich, K.; Fahrner, M.; Brombacher, E.; Seredynska, A.; Maldacker, M.; Kreutz, C.; Schmidt, A.; Schilling, O. Data-Independent Acquisition: A Milestone and Prospect in Clinical Mass Spectrometry–Based Proteomics. Molecular & Cellular Proteomics 2024, 23 (8), 100800. DOI: 10.1016/j.mcpro.2024.100800.

(27) Marino, F.; Cristobal, A.; Binai, N. A.; Bache, N.; Heck, A. J.; Mohammed, S. Characterization and usage of the EASY-spray technology as part of an online 2D SCX-RP ultra-high pressure system. Analyst 2014, 139 (24), 6520–6528. DOI: 10.1039/c4an01568a From NLM.

(28) Kovalchuk, S. I.; Jensen, O. N.; Rogowska-Wrzesinska, A. FlashPack: Fast and Simple Preparation of Ultrahigh-performance Capillary Columns for LC-MS. Mol Cell Proteomics 2019, 18 (2), 383–390. DOI: 10.1074/mcp.TIR118.000953 From NLM.

(29) Ishihama, Y.; Rappsilber, J.; Andersen, J. S.; Mann, M. Microcolumns with self-assembled particle frits for proteomics. Journal of Chromatography A 2002, 979 (1), 233–239. DOI: 10.1016/S0021-9673(02)01402-4.

(30) Maiolica, A.; Borsotti, D.; Rappsilber, J. Self-made frits for nanoscale columns in proteomics. Proteomics 2005, 5 (15), 3847–3850. DOI: 10.1002/pmic.200402010 From NLM.

(31) Demichev, V.; Messner, C. B.; Vernardis, S. I.; Lilley, K. S.; Ralser, M. DIA-NN: neural networks and interference correction enable deep proteome coverage in high throughput. Nature Methods 2020, 17 (1), 41–44. DOI: 10.1038/s41592-019-0638-x.

(32) Perez-Riverol, Y.; Bai, J.; Bandla, C.; García-Seisdedos, D.; Hewapathirana, S.; Kamatchinathan, S.; Kundu, D. J.; Prakash, A.; Frericks-Zipper, A.; Eisenacher, M.;, et al. The PRIDE database resources in 2022: a hub for mass spectrometry-based proteomics evidences. Nucleic Acids Res 2022, 50 (D1), D543–d552. DOI: 10.1093/nar/gkab1038 From NLM.

(33) Hsieh, E. J.; Bereman, M. S.; Durand, S.; Valaskovic, G. A.; MacCoss, M. J. Effects of Column and Gradient Lengths on Peak Capacity and Peptide Identification in Nanoflow LC-MS/MS of Complex Proteomic Samples. Journal of the American Society for Mass Spectrometry 2013, 24 (1), 148–153. DOI: 10.1007/s13361-012-0508-6.

(34) Köcher, T.; Swart, R.; Mechtler, K. Ultra-high-pressure RPLC hyphenated to an LTQ-Orbitrap Velos reveals a linear relation between peak capacity and number of identified peptides. Anal Chem 2011, 83 (7), 2699–2704. DOI: 10.1021/ac103243t From NLM.

(35) MacNair, J. E.; Lewis, K. C.; Jorgenson, J. W. Ultrahigh-Pressure Reversed-Phase Liquid Chromatography in Packed Capillary Columns. Analytical Chemistry 1997, 69 (6), 983–989. DOI: 10.1021/ac961094r.

(36) Stegeman, G.; Kraak, J. C.; Poppe, H. Dispersion in packed-column hydrodynamic chromatography. Journal of Chromatography A 1993, 634 (2), 149–159. DOI: 10.1016/0021-9673(93)83001-9.

(37) Lauber, M. A.; Koza, S. M.; Fountain, K. J. Optimizing Peak Capacity in Nanoscale Trap-Elute Peptide Separations with Differential Column Heating. 2014, Waters. 720005047EN.

(38) Xu, P.; Duong, D. M.; Peng, J. Systematical Optimization of Reverse-Phase Chromatography for Shotgun Proteomics. Journal of Proteome Research 2009, 8 (8), 3944–3950. DOI: 10.1021/pr900251d.

(39) Wang, H.; Yang, Y.; Li, Y.; Bai, B.; Wang, X.; Tan, H.; Liu, T.; Beach, T. G.; Peng, J.; Wu, Z. Systematic Optimization of Long Gradient Chromatography Mass Spectrometry for Deep Analysis of Brain Proteome. Journal of Proteome Research 2015, 14 (2), 829–838. DOI: 10.1021/pr500882h.

(40) Gritti, F.; Guiochon, G. Overload behavior and apparent efficiencies in chromatography. Journal of Chromatography A 2012, 1254, 30–42. DOI: 10.1016/j.chroma.2012.07.015.

## References

(1) Kovalchuk, S. I.; Jensen, O. N.; Rogowska-Wrzesinska, A. FlashPack: Fast and Simple Preparation of Ultrahigh-performance Capillary Columns for LC-MS. Mol Cell Proteomics 2019, 18 (2), 383–390. DOI: 10.1074/mcp.TIR118.000953 From NLM.

(2) Ishihama, Y.; Rappsilber, J.; Andersen, J. S.; Mann, M. Microcolumns with self-assembled particle frits for proteomics. Journal of Chromatography A 2002, 979 (1), 233–239. DOI: 10.1016/S0021-9673(02)01402-4.

(3) Maiolica, A.; Borsotti, D.; Rappsilber, J. Self-made frits for nanoscale columns in proteomics. Proteomics 2005, 5 (15), 3847–3850. DOI: 10.1002/pmic.200402010 From NLM.

(4) Demichev, V.; Messner, C. B.; Vernardis, S. I.; Lilley, K. S.; Ralser, M. DIA-NN: neural networks and interference correction enable deep proteome coverage in high throughput. Nature Methods 2020, 17 (1), 41–44. DOI: 10.1038/s41592-019-0638-x.

(5) Perez-Riverol, Y.; Bai, J.; Bandla, C.; García-Seisdedos, D.; Hewapathirana, S.; Kamatchinathan, S.; Kundu, D. J.; Prakash, A.; Frericks-Zipper, A.; Eisenacher, M.;, et al. The PRIDE database resources in 2022: a hub for mass spectrometry-based proteomics evidences. Nucleic Acids Res 2022, 50 (D1), D543–d552. DOI: 10.1093/nar/gkab1038 From NLM.

